# Interactive Analysis of Biosurfactants in Fruit-Waste Fermentation Samples using BioSurfDB and MEGAN

**DOI:** 10.1101/2021.11.11.468240

**Authors:** Gabriela Fiori da Silva, Anupam Gautam, Iolanda Cristina Silveira Duarte, Tiago Palladino Delforno, Valéria Maia de Oliveira, Daniel H. Huson

## Abstract

Microbial biosurfactants are of major interest due to their multifunctional properties, biodegradable nature and low toxicity. Agroindustrial waste, such as fruit waste, can be used as substrates for producing bacteria. In this study, six samples of fruit waste, from oranges, mangoes and mixed fruits, were self-fermented, and then subjected to short-read metagenomic sequencing, so as to allow assessment of the potential of the associated microbiota for biosurfactant production. Taxonomic analysis using the DIAMOND+MEGAN analysis pipeline shows that all six samples are dominated by Proteobacteria, in particular, a common core consisting of the genera *Klebsiella*, *Enterobacter*, *Stenotrophomonas*, *Acinetobacter* and *Escherichia*. To support the interactive visualization and exploration of the surfactant-related genes in such samples, we have integrated the BiosurfDB classification into MEGAN and make this available. Functional analysis indicates high similarity among samples and that a significant number of reads map to genes that are involved in the biosynthesis of lipopeptide-class biosurfactants. Gene-centric analysis reveals *Klebsiella* as the main assignment for genes related to putisolvins biosynthesis. This suggests that fruit waste is a promising substrate for fermentative processes because the associated microbiota may be able to produce biosurfactants that are potentially useful for the agricultural, chemical, food and pharmaceutical industries.

## Introduction

Biosurfactants are surface-active molecules produced by microorganisms that have been highlighted as an environmentally-friendly alternative to their synthetic counterpart, chemical surfactants^1,2^, which are produced by the petrochemical industry. Microbial surfactants demonstrate higher degradability, lower toxicity, selectivity, antimicrobial and anti-adhesive properties and applicability in large pH, temperature, and salinity spectra^3–5^. They are versatile and have applications in pharmaceutics, food, cosmetics, agriculture, wastewater, bioremediation, enhanced oil recovery, metal removal and other industrial sectors^6–8^.

Biosurfactant-producing microorganisms have been reported and isolated from several sources such as marine habitats, mangroves, freshwater, soil, sludge and fruits^9–12^. Fruit waste and residues generated by fruit processing industries present a renewable and low-cost carbon source for fermentation in biosurfactant production^12–14^. To allow better exploitation of such waste as a source of useful microorganisms, a clearer understanding of the taxonomic and functional diversity of the present microbes is required. Shotgun metagenomic sequencing and subsequent alignment-based analysis allows one to identify specific genes of interest that are related to biosurfactant production.

Various metagenomic approaches combined with bioinformatics tools have been used to study biosurfactants^15^. However, general-purpose functional databases, such as KEGG^16^, are not ideal for this purpose as they tend to focus on other aspects of function. In particular, some genes related to biosurfactant biosynthesis are classified under antibiotics in nonribosomal peptide pathways. Hence, the use of a domain-specific database is indicated^17^.

Here we present six short-read metagenomic datasets that we have collected from different fruit-fermentation batch reactors using short-read Illumina sequencing. We used the DIAMOND+MEGAN^18^ pipeline to determine the microbial diversity of the fruit-waste samples and to investigate the functional potential for biosurfactant production. In more detail, taxonomy analysis was performed by MEGAN based on DIAMOND alignments against the NCBI-nr protein reference database^19^. For functional analysis, here we present a new extension of MEGAN that allows functional analysis based on the BiosurfDB database, a domain-specific database focused on identifying genes related to biosurfactant production and biodegradation^20^.

Our analysis indicates that the six samples possess a common core of Gammaproteobacteria, dominated by the genera *Klebsiella*, *Enterobacter*, *Stenotrophomonas*, *Escherichia* and *Acinetobacter*. The samples have similar functional profiles that show a potential for biosurfactant biosynthesis, especially lipopeptides.

To firmly establish the taxonomic identity of the organisms that contain the genes related to biosurfactant production, we applied gene-centric assembly^21^ to those genes and aligned the resulting gene-length contigs against the NCBI-nt database^19^ using BLASTN^22^.

## Results

### Taxonomic analysis

Each of the six metagenomic datasets where sequenced to over 20 million reads per sample, using Illumina sequencing in a 2 × 150 bp layout. Of 140, 183, 232 total input sequencing reads, 121, 875, 304 obtained at least one alignment against the NCBI-nr database, and of these, 121, 561, 304 could be assigned to a taxon in the NCBI taxonomy, leading to an assignment rate of 87%.

In all samples, reads were predominantly assigned to bacteria by an NCBI-nr run of the DIAMOND+MEGAN pipeline. The three main phyla are Proteobacteria followed by Firmicutes and Actinobacteria (Table 1). Rarefaction analysis (see Supplementary Figure S1) indicated that the number of genera detected reached a plateau for each of the samples, indicating that the amount of sequencing performed was sufficient to obtain a stable representation of the taxonomic content of the samples. Rarefaction analysis suggests that the Mango and Mix samples have slightly higher diversity than the Orange samples, which is also reflected in the Shannon-Weaver diversity indices calculated from the genus-level assignments, which are 2.1, 1.5, 2.0, 1.8, 0.5 and 1.0, for the the Mango-1, Mango-2, Mix-1, Mix-2, Orange-1 and Orange-2 samples, respectively.

**Table 1.** For each of the six fruit samples, we report the total percentage of reads assigned the bacterial domain and to three top bacterial phyla.

|  | Mango-1 | Mango-2 | Mix-1 | Mix-2 | Orange-1 | Orange-2 |
| --- | --- | --- | --- | --- | --- | --- |
| **Domain** |  |  |  |  |  |  |
| Bacteria | 99.9 | 99.9 | 99.9 | 99.9 | 99.8 | 99.9 |
| **Phylum** |  |  |  |  |  |  |
| Proteobacteria | 99.1 | 98.3 | 95.8 | 96.8 | 99.0 | 99.6 |
| Firmicutes | 0.6 | 1.3 | 4.0 | 3.1 | 1.0 | 0.4 |
| Actinobacteria | 0.2 | 0.4 | 0.1 | 0.1 | 0.0 | 0.0 |

Most of the taxonomic content of the six samples falls into 19 genera (Figure 1). *Klebsiella* dominates all six samples, accounting for ≈ 92 % and 84 % of genus-level assignments in the Orange-1 and Orange-2 samples respectively, and with assignment rates of ≈ 32 %, 69 %, 44 % and 49 %, for Mango-1, Mango-2, Mix-1 and Mix-2, respectively. Differences in the proportion of reads assigned to specific genera were observed between all six samples, which is to be expected even between duplicates, as the duplicates are biological replicates (Table 2).

**Table 2.**
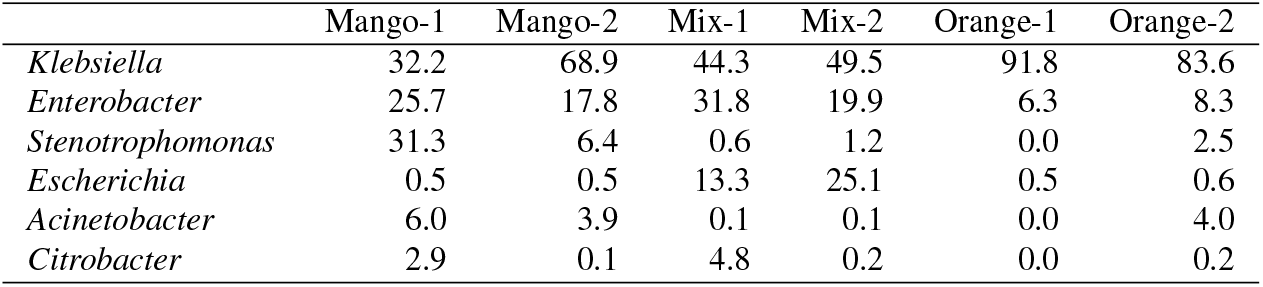
For each of the six fruit samples, we report the we report the relative percentage of reads assigned to the 6 top bacterial genera.

**Figure 1.**
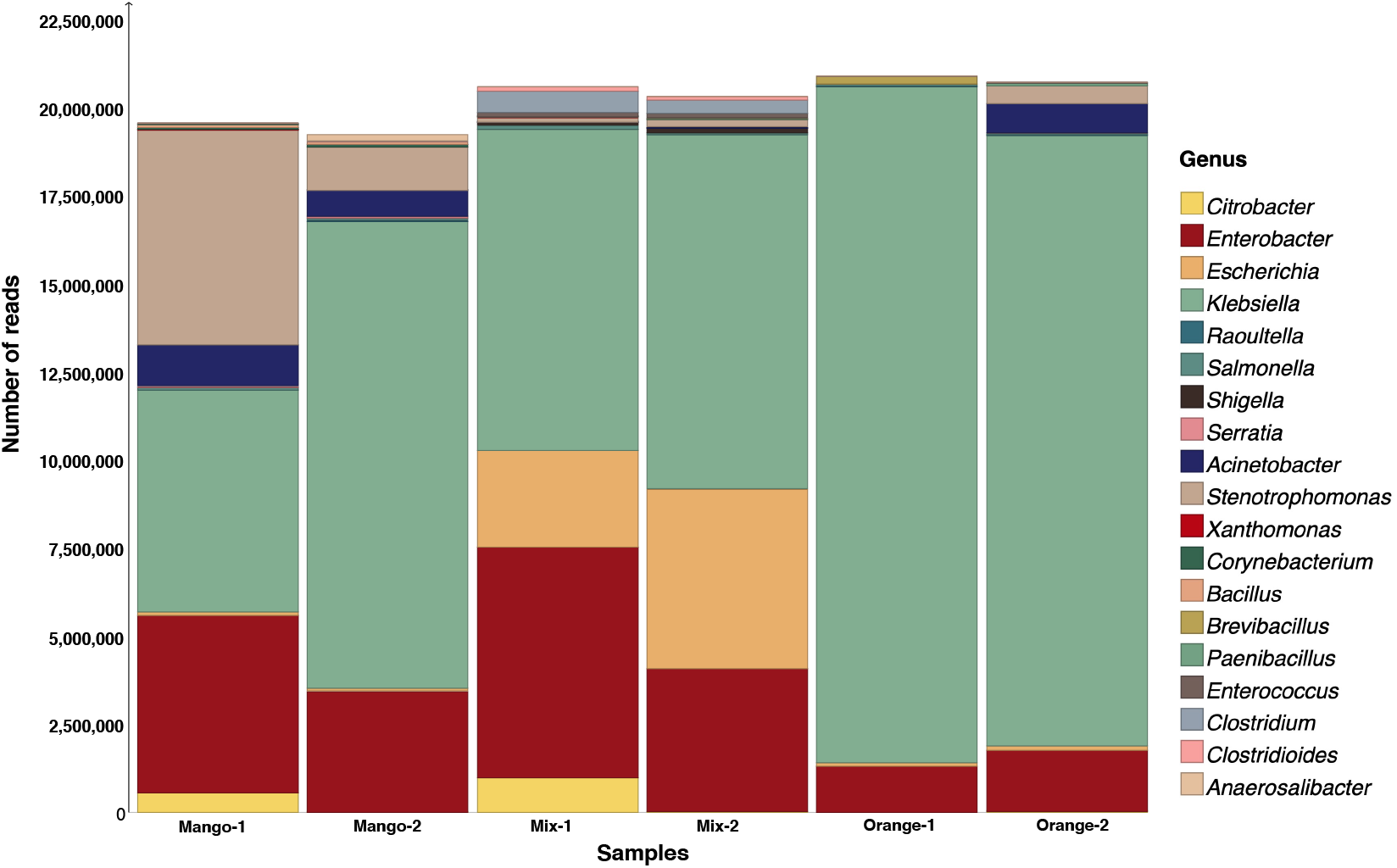
For each of the six fruit samples, we show the number of reads assigned to each of 19 occurring genera, based on an NCBI-nr run of the DIAMOND+MEGAN pipeline. Note that “taxonomic projection” was performed to map all read assignments to the genus level.

The NCBI-nr runs of MEGAN indicate that the six fruit samples studied here are dominated by *Gammaproteobacteria*, with most reads assigned to the genera *Klebsiella* (*≈* 63 %), *Enterobacter* (*≈* 19 %), *Stenotrophomonas* (*≈* 7 %), *Escherichia* (≈ 7 %) and *Acinetobacter* (≈ 2 %). The first, second and fourth genera are members of the Enterobacteriaceae family, whereas the third and last belong to the family of Lysobacteriaceae and Moraxellaceae, respectively.

### Functional analysis

The results of the BioSurfDB runs of the DIAMOND+MEGAN pipeline were loaded into MEGAN. This provided an overview of the functional potential present in the samples and allowing an assessment of the relative abundances of genes related to biosynthesis of biosurfactants (Figure 2). Strikingly, more than 60% of all reads that map to a Surfactant class were assigned to a biosynthesis of lipopeptide biosurfactant subclass. In addition, a fair number of reads were aligned to genes crucial for synthesizing non-ribosomal lipopeptide structures, namely surfactin, mycosubtilin, iturin A, lichenysin D, bacillomycin D and fengycin (Figure 3).

**Figure 2.**
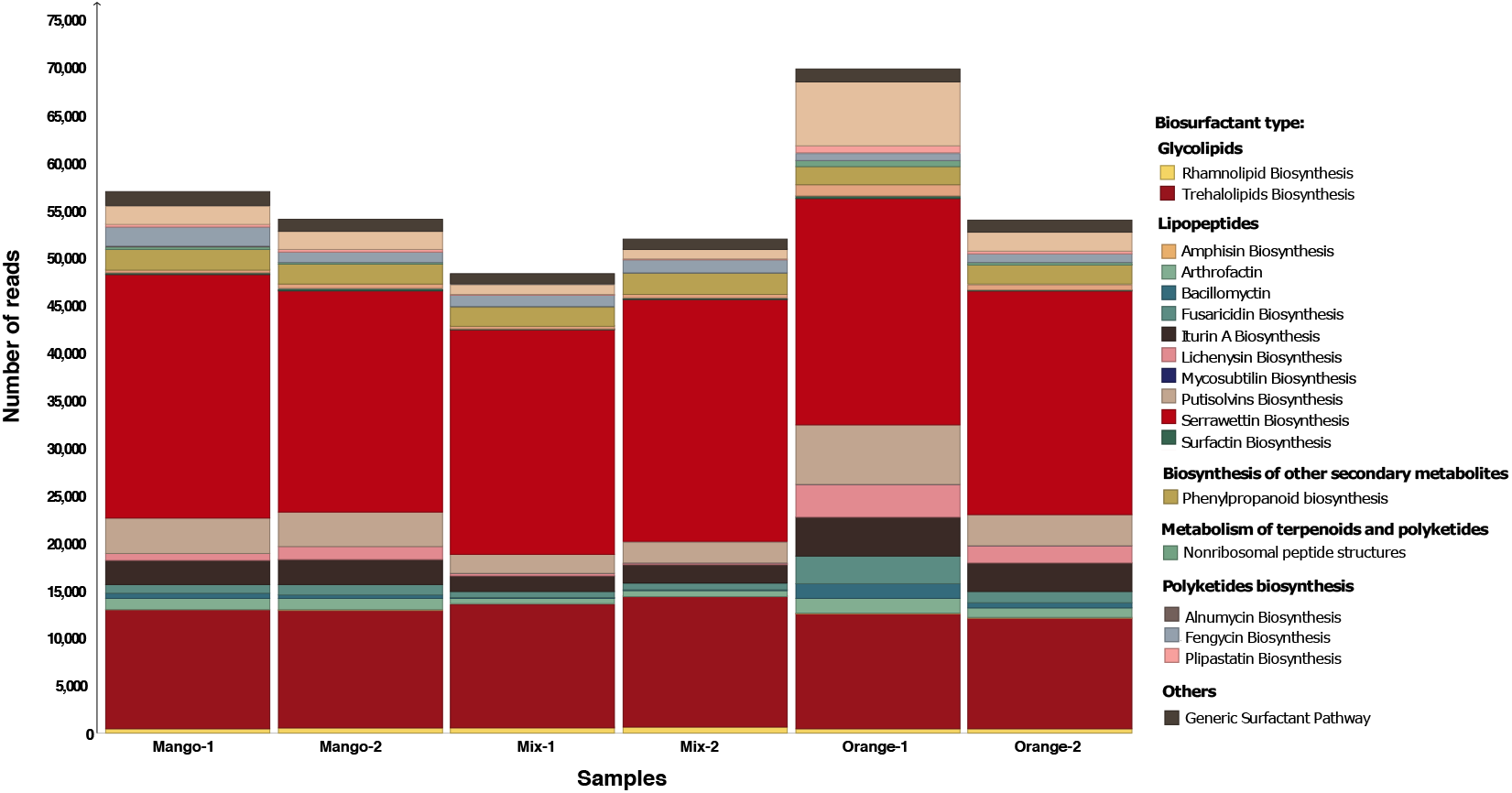
For each of the six fruit samples, we report the number of reads that are assigned to a surfactant class, based on a BioSurfDB run of the DIAMOND+MEGAN pipeline.

**Figure 3.**
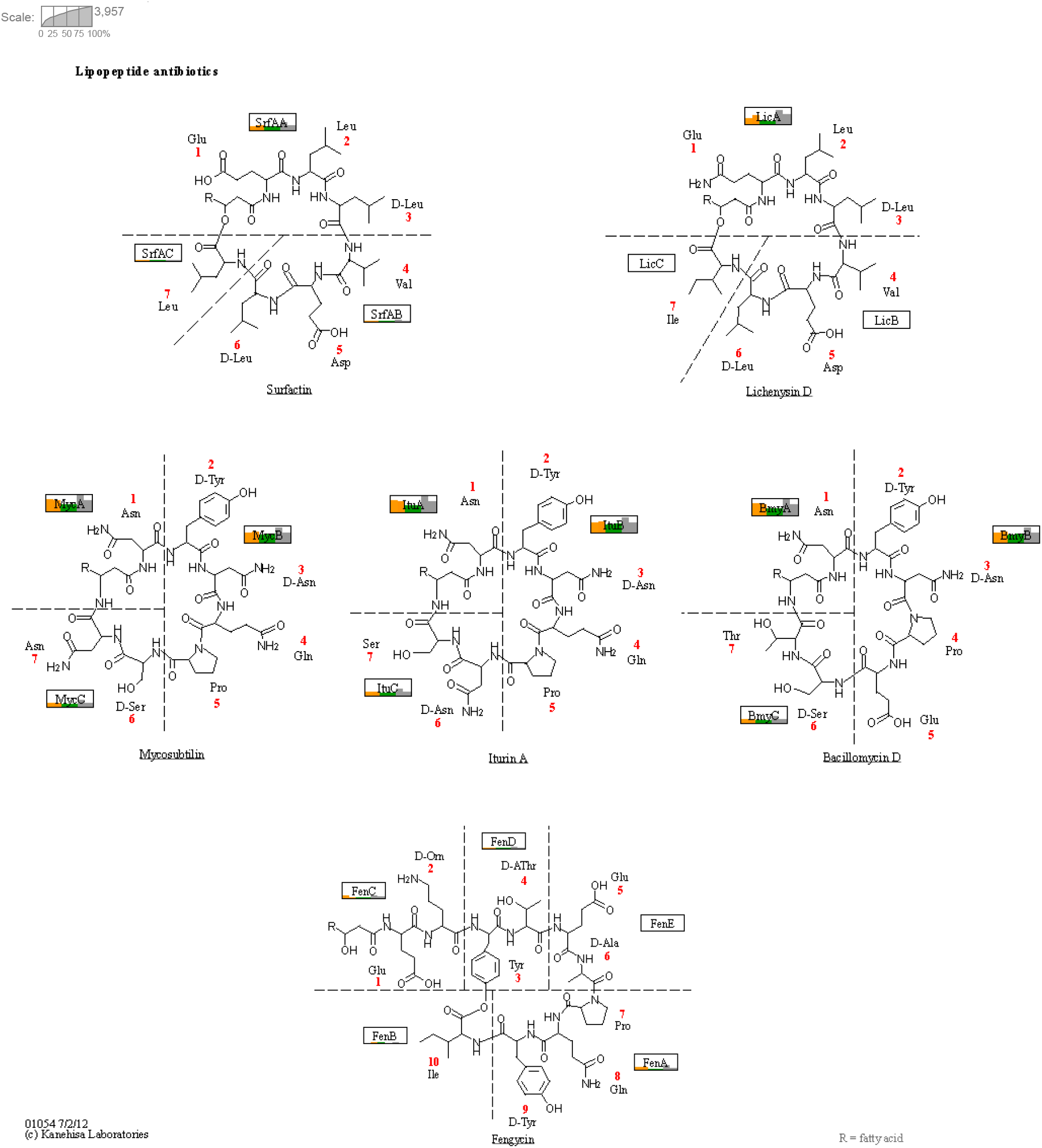
For each of the six fruit-based samples, we indicate the number of reads assigned to the different genes involved in non-ribosomal synthesis. The illustration is based on KEGG pathway ko01054 “Nonribosomal peptide structures”^16^. Each gene is represented by a rectangle and inside each rectangle we use a bar chart to indicate the number of reads assigned from each of the six samples, namely Mango-1 and Mango-2 (yellow), Mix-1 and Mix2 (green) and Orange-1 and Orange-2 (gray).

In more detail, the most abundant proteins present in the samples were related to the biosynthesis of pultisolvins followed by trehalolipids, mycosubtilin and iturin A. Trehalolipids are classified as glycolipids while putisolvins, mycosubtilin and iturin A are considered lipopeptides. Significant differences were observed only in the lipopeptide biosynthesis category: iturin A biosynthesis (Figure 4a) was more abundant in Orange samples than Mix and Mango samples and amphisin biosynthesis (Figure 4b) was more abundant in Orange samples than in Mix samples.

**Figure 4.**
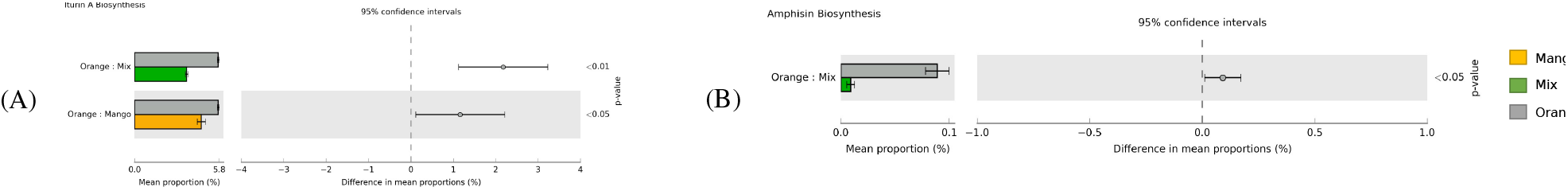
Comparison of pairs of samples grouped by type of fruit against each other, such as Orange vs. Mix. Here we show two biosurfactant types whose mean proportion of assigned reads exhibit significant differences (p < 0.05), namely (A) Iturin A Biosynthesis and (B) Amphisin Biosynthesis.

To address the question of which taxa contain the detected biosurfactant genes, we applied gene-centric assembly to reads assigned to a number of such genes and then determined the taxonomic assignment of the resulting contigs. For a cluster of genes involved in putisolvins production, the resulting contigs were mainly assigned to genera in the above described common core of genera found in the fruit samples. The class Gammaproteobacteria, the family Enterobacteriaceae, and their representatives *Klebsiella* sp. and *Enterobacter* sp. (Figure 5) were most assigned. At a lower abundance, *Acinetobacter* sp. and *Stenotrophomonas* sp. were also assigned.

**Figure 5.**
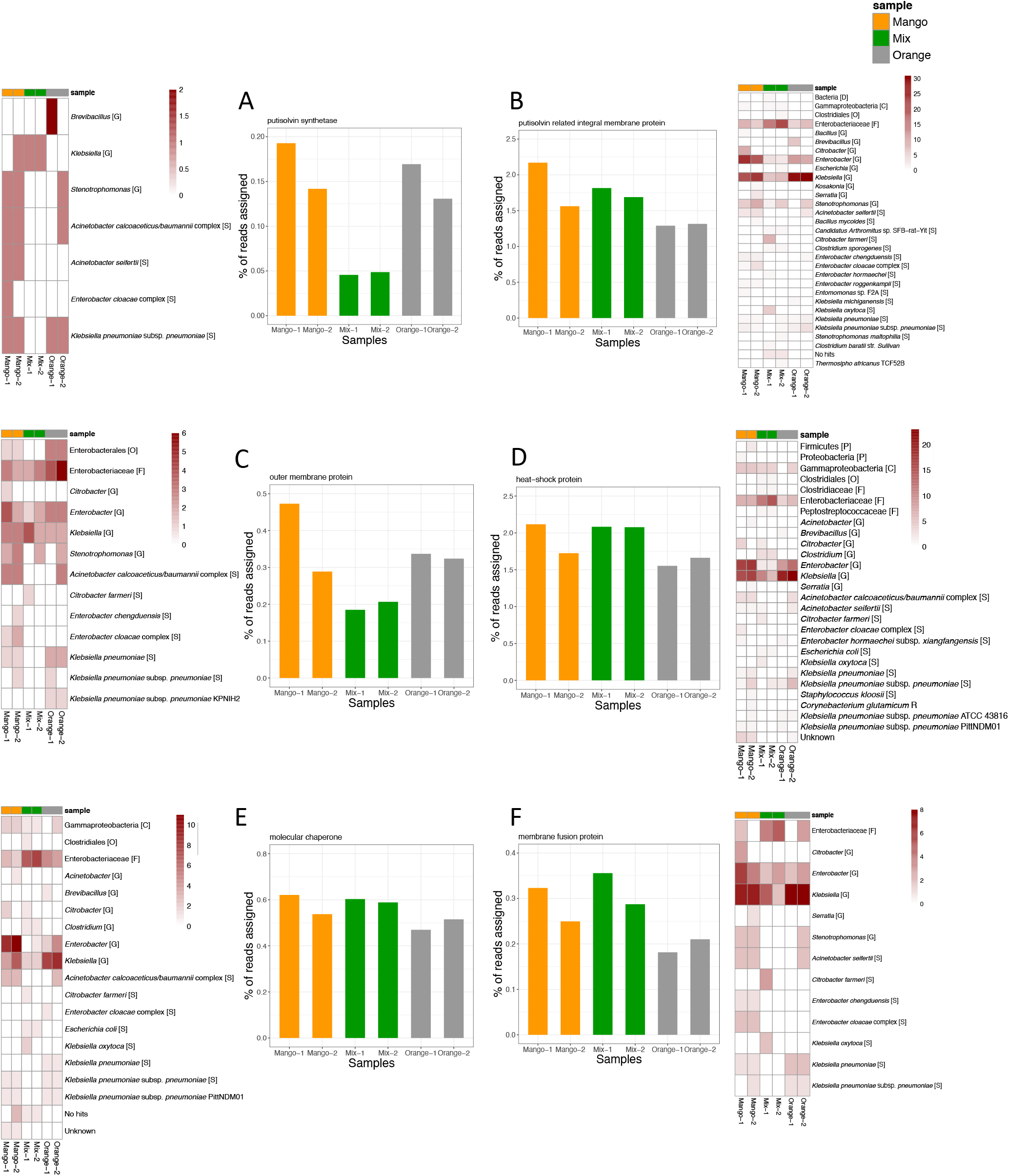
For each six different genes involved in putisolvins biosynthesis, we report under (A) - (F) the percentage of reads assigned to the gene, for each sample. For each gene, we also show a heatmap indicating how many contigs obtained by gene-centric assembly where assigned to certain taxa, based on their alignment against the NCBI-nt database.

The taxonomic assignment for contigs obtained from genes involved in the synthesis of mycosubtilin, iturin A and lichenysin (Figure 3) were mainly from classes and families associated with the mentioned common core of genera. The genera *Brevibacillus*, *Klebsiella* and *Enterobacter* received the highest assignments (Figure S2).

These results suggest that all six waste fruit samples have a similar functional profile related to biosurfactant biosynthesis and all samples appear to have a significant potential for the production of biosurfactants.

## Discussion

Our analysis suggests that the microbiota of the six investigated fruit samples share a common core of Gammaproteobacteria and that members of the family Enterobacteriaceae are the most abundant. This result is compatible with previous studies in which Enterobacteriaceae were shown to be predominant in plant-associated microbiomes^23,24^. Some genera are commonly reported as endophytic bacteria, such as *Enterobacter* and *Klebsiella*, which can play an important role in plant survival, production of toxins, antimicrobial compounds, and even contribute to plant growth promotion^25^. However, in general, fruit microbiota are diverse and composition are influcenced by management practices, pesticide use, external factors, growing season, etc.^26^.

Several genera of Proteobacteria are considered efficient oil-degrading and biosurfactant-producing microorganisms, such as *Pseudomonas*, *Streptomyces*, *Enterobacter*, *Acinetobacter*, *Escherichia*, *Klebsiella*, and *Stenotrophomonas*^3,27–31^. Some of these appear as members of the microbial common core presented here.

Our study indicates that the functional profiles of the microbiota present in the six fruit samples are quite similar to each other and have the potential for biosurfactant biosynthesis, especially of lipopeptides. Lipopeptides biosurfactants are considered potent antimicrobials that are capable of disrupting the cell membrane^32^. In addition, they also exhibit antitumor, immunomodulatory, emulsifying activities and their production is also closely related to motility and biofilm formation or inhibition^15,33,34^. These biosurfactants are synthesized by a set of multifunctional enzymes called non-ribosomal peptide synthases (NRPSs). They can have diverse structures, and nutritional parameters can influence their composition^35,36^. The high occurrence of genes related to the biosynthesis of lipopeptide biosurfactants may be linked to the fact that some endophytic bacteria play a protective role to plants against pathogens and produce such antimicrobial molecules^36,37^.

Previous studies have reported that putisolvins biosurfactants are closely related to motility, in the dispersion of naphthalene and phenanthrene crystals and also in breaking or inhibiting biofilms^38^. Their production depends on several regulatory genes such as the heat shock system DnaK and the two component system GacA/GacS^39^. Their production has has been mainly attributed to the *Pseudomonas putida* species^38,39^, whereas our study links them to *Klebsiella*, *Enterobacter* and *Acinetobacter*. This taxonomic assignment highlights the presence of certain regulatory genes for secondary metabolites common to gram-negative bacteria^39^. Furthermore, the knowledge that these bacteria possess genes involved in the biosynthesis of putisolvins may direct studies on the production of these molecules from these microorganisms and theirs applicability in biofilm removal.

Another type of biosurfactant present in our samples is iturin A and this specific class of biosurfactants, iturins, can have different conformations due to amino acid variations, and its main variants are: iturin A, iturin C, mycosubtilin, bacillomycin D, F, L and LC^1,37^. *Brevibacillus*, *Klebsiella* and *Enterobacter* were the genera for which the most reads are assigned to genes related to the biosynthesis of the iturins family. In addition, all fruit samples have some reads that align to genes involved in the synthesis of such molecules, namely srfAA, licA, ituA and BmyA that are related to surfactin, lichenysin. mycosubitilin, iturin A and bacillomycin respectively, and the abundance of these This suggests that these microbiota are a promising source of these molecules.

Biosurfactants mainly arise in response to environmental conditions imposed on the microorganism, acting in physiological functions such as motility, protection from toxins, adhesion to substrates and cell interactions^40^. The microbiota of hydrocarbon-contaminated sites have been widely studied, isolated and used in the production of biosurfactants due to the ability of these microorganisms to utilize hydrophobic carbon sources and thus play a role in bioremediation. This work provides a new perspective for the prospection of biosurfactant-producing microorganisms, as it suggests that fruit-waste samples are a promising source of bacteria capable of producing biosurfactants, mainly lipopeptides, which may have application in several industrial sectors. Furthermore, isolation of culture-dependent strains from these fruit residues might be usable in fermentative processes, and we envision testing of various nutritional and temperature parameters can be evaluated for the production of these medically and industrially important molecules.

Our analysis determined similar profiles directed towards biosurfactant biosynthesis. The samples show high reads counts for biosurfactant production, mainly lipopeptides, a potential source of novel antibiotic and antifungal molecules. Furthermore, in line with the result of taxonomic analysis, the results of the gene-centric analysis showed that the *common core* found in the samples is directly related to the genes of interest.

The BiosurfDB domain-specific database is an essential resource for detecting the presence of genes associated biosurfactant biosynthesis. Integration of the BiosurfDB classification into our interactive metagenome analysis tool allows the user to interactively explore and compare the reads assigned to biosurfactant genes.

While the putisolvins genes detected in our samples are represented in BioSurfDB by reference genes obtained from *Pseudomonas* genomes, gene-centric assembly and DNA alignment to the NCBI-nt database clearly show that the organisms carrying these genes in the six samples are not *Pseudomonas*, but rather belong to the genera *Klebsiella*, *Enterobacter* and *Acinetobacter*.

## Methods

### Sampling and autochthonous fermentation

Fruit residues were collected from open fairs in the city of Sorocaba, São Paulo, Brazil. The fruit residues were separated into three distinct samples: orange bagasse, mango residue and mixed fruit residue (using a mixture of fruits that includes papaya, pear, avocado, grapes, guava and banana residues). Each fruit sample was crushed and 25 g L^*−*1^ of total solids was added to a batch reactor with Luria-Bertami (LB) medium, in duplicate. In all six batch reactors, fermentation assays were conducted for 5 days at 32 °C and 150 rpm.

### DNA extraction and metagenome sequencing

DNA from each autochthonous fermentation was extracted and purified using the PowerSoil DNA kit (MoBio Laboratories, Inc., Carlsbad, CA, USA), following the manufacturer’s instructions. DNA concentration and quality were estimated using a ND-2000 spectrophotometer (Nanodrop Inc, Wilmington, DE), using a ratio of 260/280 nm > 1.8. For each extraction, the DNA was sequenced in a 2 × 150 bp format using a Illumina HiSeq at the Animal Biotechnology Laboratory, Department of Animal Science (ESALQ/USP, Piracicaba, São Paulo, Brazil) according to the manufacturer’s guidelines. This resulted in six datasets, which we will refer to as Orange-1, Orange-2, Mango-1, Mango-2, Mix-1 and Mix-2. These contain very similar numbers of reads, namely 23407430, 22989500, 24136468, 24072864, 21872692, and 23, 704, 278, respectively.

All sequencing reads were submitted to the European Nucleotide Archive under project accession PRJEB47062 and sample accessions ERS7265231 (Mango-1), ERS7265232 (Mango-2), ERS7265233 (Mix-1), ERS7265234 (Mix-2), ERS7265235 (Orange-1) and ERS7265236 (Orange-2).

### BioSurfDB representation in MEGAN

The MEGAN software allows incorporation of additional classifications. Here we describe how to add a new functional classification to MEGAN that represents data provided by the BioSurfDB^20^ database, which is focused on biosurfactants and biodegradation. Using the URLs https://www.biosurfdb.org/api/get/pathway, https://www.biosurfdb.org/api/get/protein and https://www.biosurfdb.org/api/get/organism_pathway, we downloaded three files in JSON format from BioSurfDB, which we called `pathway.json`, `protein.json` and `organism_pathway.json`, respectively.

The first file, `pathway.json`, contains information about pathways. We parsed it to create the basic hierarchical tree representation of entities in the BioSurfDB classification. This representation uses integer identifiers for all nodes in the tree and a separate mapping of those identifiers to names. We provide the tree representation and the mapping in two files called `biosurfdb.tre` and `biosurfdb.map`, respectively.

The second downloaded file, `protein.json`, provides all protein accessions and sequences that occur in the classification. We provide these in a fasta file called `biosurfdb.fasta`. Finally, the third downloaded file, `organism pathway.json`, was used to place the proteins into the tree representation. As the number of proteins is small (≈ 6,000), their accessions were also added to the tree as leaves. The downloaded file was also used to create a mapping of protein accessions to the integer node identifiers. We provide this mapping as the file `acc2biosurfdb.map`.

As mentioned, MEGAN allows the user to incorporate new classifications into the software. This requires that one prepares two files that describe the hierarchical structure of the classification, such as the two files `biosurfdb.tre` and `biosurfdb.map` described above. Use the `Edit-Preferences-Add Classification`… menu item to select the tree file so as to add the new classification to MEGAN. The program will provide a viewer for the classification and will integrate it into all menus, toolbars and dialogs.

To enable MEGAN to assign reads to a new classification, during the meganization or import of an alignment file, the user must specify an appropriate mapping file that maps reference sequence accessions to entities in the classification. In the case of alignment against the BioSurfDB sequences in `biosurfdb.fasta`, the appropriate mapping file is `acc2biosurfdb.map`.

### DIAMOND+MEGAN analysis

All six samples were first aligned against the NCBI-nr database^19^ (downloaded from ftp://ftp.ncbi.nih.gov/blast/db/FASTA/nr.gz in January 2021) using DIAMOND^41^ (version 2.0.0, default alignment options). The resulting DAA files were “meganized”, i.e. subjected to taxonomic and functional binning, using MEGAN^18^ (version 6.21.2, default options). We will refer to this DIAMOND+MEGAN analysis as an NCBI-nr run.

We also aligned all six datasets against the BioSurfDB reference file using DIAMOND (default alignment options), The resulting DAA files were “meganized” with the help of the BioSurfDB-specific mapping file `acc2biosurf.map`. We will refer to ths DIAMOND+MEGAN analysis as an BioSurfDB run.

Taxonomic and functional profiles were exported from MEGAN and statistical analysis was performed using STAMP^42^. ANOVA was conducted and Tukey’s adjustment was applied to determine the significant differences between samples. Unclassified reads were removed from the analysis and results with *p* < 0.05 (corrected *p*-value) were considered significant.

To determine the genus-level taxonomic context of the samples, we employed *taxonomic projection* as implemented in MEGAN. In this calculation, all reads assigned at a taxonomic rank that is more specific than genus are “projected up” to the appropriate genus, whereas all reads that are assigned at a higher taxonomic rank are “projected down” onto the subsequent genera, in proportion to the number of reads assigned to each such genus.

### Gene-centric assembly

MEGAN provides an algorithm for assembling all reads that align to a specific class of reference sequences, using protein-alignment-guided assembly, as described in^21^. For each of the six samples, we applied this algorithm (default parameters) to genes associated with Surfactant classes in the BioSurfDB classification.

For the Putisolvins Biosynthesis class we assembled the genes for heat-shock protein, membrane fusion protein, molecular chaperone, outer membrane protein, putisolvin related integral membrane protein, and putisolvin synthetase (see Table 3). For the classes Iturin A Biosynthesis, Lichenysin Biosynthesis and Mycosubtilin Biosynthesis, we assembled the genes for Iturin A synthetase C, lichenysin synthetase A, and both mycosubtilin synthase subunit A and Mycosubtilin synthase subunit B, respectively.

**Table 3.** Basic statistics for the gene-centric assembly of surfactant-related genes. For each of six genes associated with the class Putisolvins Biosynthesis, we report the number of contigs, their minimum, mean and maximum length and their average coverage. The values reported here are for the Mango-1 sample, the values for the other samples are similar and are reported in the Supplement.

| Gene name | Number of contigs | Min. | Mean length | Max. | Average coverage |
| --- | --- | --- | --- | --- | --- |
| Heat-shock protein | 64 | 201 | 575 | 1572 | 7.2 |
| Membrane fusion protein | 26 | 207 | 257 | 330 | 5.0 |
| Molecular chaperone | 26 | 225 | 390 | 528 | 8.0 |
| Putisolvin related integral membrane protein | 84 | 201 | 377 | 750 | 6.4 |
| Putisolvin synthetase | 5 | 201 | 245 | 333 | 4.3 |
| Outer membrane protein | 19 | 204 | 254 | 519 | 6.1 |

The resulting contigs were aligned against the NCBI-nt database (downloaded October, 2021) using BLASTN^22^ (default parameters) and the resulting files were then imported into MEGAN so as to perform taxonomic analysis (default parameters).

## Data Availability

The six sequences are available from the European Nucleotide Archive under project accession PRJEB47062 and sample accessions ERS7265231 (Mango-1), ERS7265232 (Mango-2), ERS7265233 (Mix-1), ERS7265234 (Mix-2), ERS7265235 (Orange-1) and ERS7265236 (Orange-2).

The BioSurfDB extension for MEGAN is available here: https://software-ab.informatik.uni-tuebingen.de/download/megan6/biosurfdb.zip.

The MEGAN files for all six samples, and the results of gene-centric assembly, are available here: https://software-ab.informatik.uni-tuebingen.de/download/public/surfactant.

## Acknowledgements

We acknowledge hardware support by the High Performance and Cloud Computing Group at the Zentrum für Datenverarbeitung of the University of Tübingen, the state of Baden-Württemberg through bwHPC, the German Research Foundation (DFG) through grant no. INST 37/935-1 FUGG. We also acknowledge support of the BMBF-funded de.NBI Cloud within the German Network for Bioinformatics Infrastructure (de.NBI) (031A532B, 031A533A, 031A533B, 031A534A, 031A535A, 031A537A, 031A537B, 031A537C, 031A537D, 031A538A). This work was also supported by Coordenação de Aperfeiçoamento de Pessoal de Nível Superior - Brasil (CAPES) - finance code 001.

The authors acknowledge infrastructural support by the cluster of Excellence EXC2124 Controlling Microbes to Fight Infection (CMFI), project ID 390838134.

Furthermore, we acknowledge support by the Open Access Publishing Fund of University of Tübingen.

## Author Contribution

GFS, ICSD, TPD, and VMO conceived and planned the experiments. GFS, AG and DHH conceived and carried out the bioinformatics analysis. GFS, AG and DHH wrote the manuscript. All authors provided critical feedback and helped shape the research, analysis and manuscript.

