## Supplementary material for "Interactive Analysis of Biosurfactants in Fruit-Waste Fermentation Samples using BioSurfDB and MEGAN"

### Supplementary Data

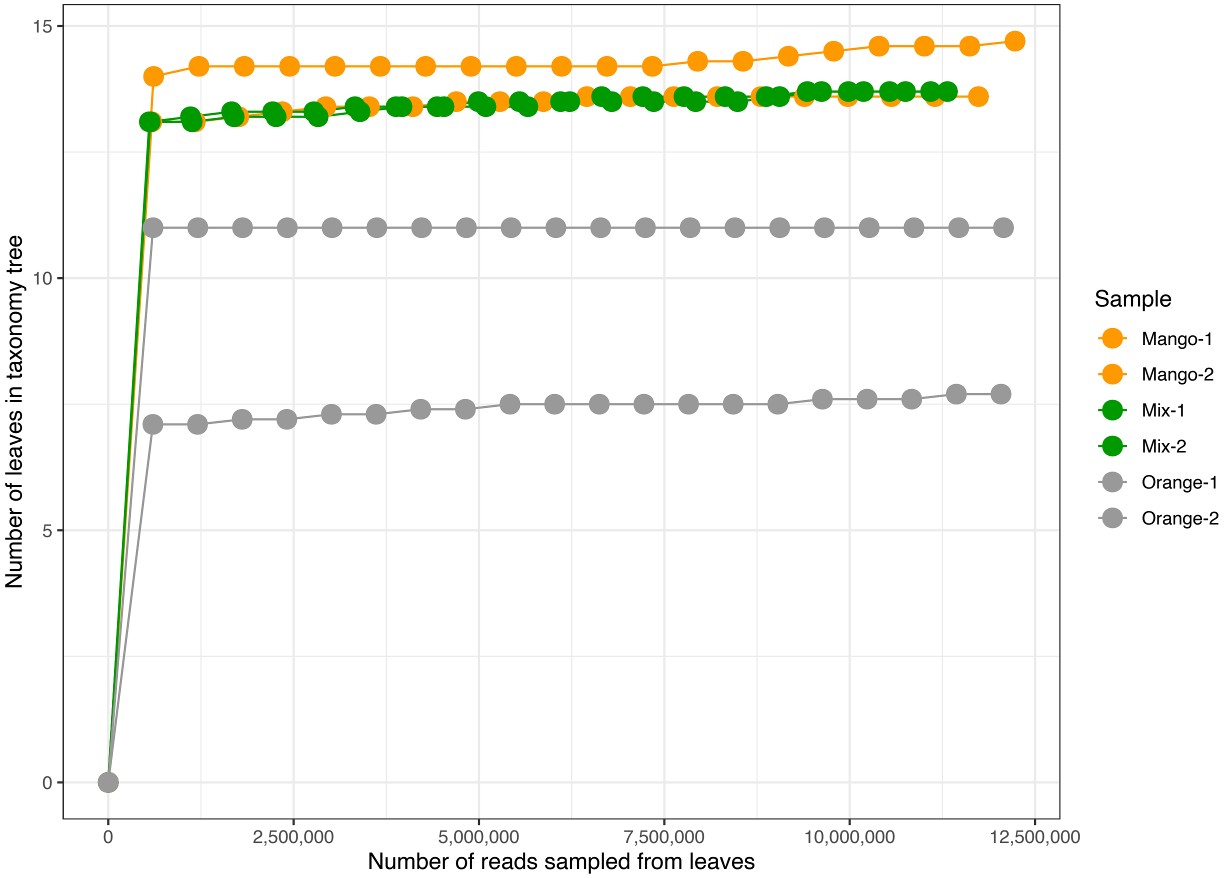


Figure 1: Rarefaction curves for six metagenome sequence data showing genera diversity over number of reads.

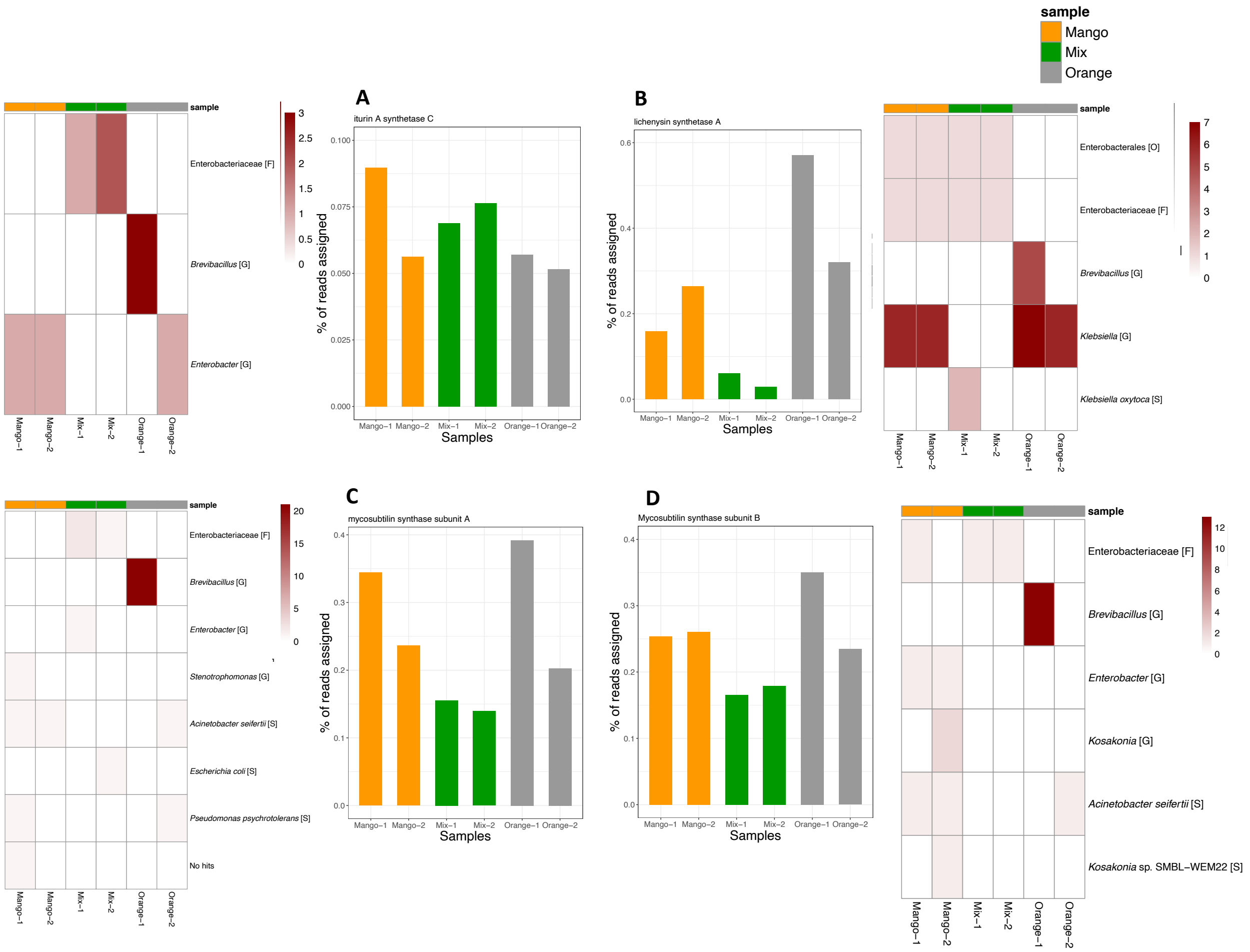


Figure 2: For one gene involved in iturin A biosynthesis, one in lichenysin biosynthesis and two in mycosubtilin biosynthesis we report under (A) - (D) the percentage of reads assigned to the gene, for each sample. For each gene, we also show a heatmap indicating how many contigs obtained by gene-centric assembly where assigned to certain taxa, based on their alignment against the NCBI-nt database.

Table 1: Basic statistics for the gene-centric assembly of surfactant-related genes. For each of six genes associated with the class Putisolvins Biosynthesis, we report the number of contigs, their minimum, mean and maximum length and their average coverage, based on the separate assembly of all six samples.

| Gene name | Number of contigs | Min. | Mean Length | Max. | Average coverage |
|---|---|---|---|---|---|
| **Mango-2** | | | | | |
| Heat-shock protein | 68 | 201 | 561.2 | 1110 | 6.5 |
| Membrane fusion protein | 21 | 216 | 264.1 | 321 | 5.1 |
| Molecular chaperone | 32 | 207 | 359.4 | 528 | 6.7 |
| Outer membrane protein | 19 | 204 | 258.9 | 519 | 6.2 |
| Putisolvin related integral membrane protein | 78 | 201 | 387.3 | 777 | 5.9 |
| Putisolvin synthetase | 5 | 210 | 276.6 | 333 | 3.8 |
| **Mix-1** | | | | | |
| Heat-shock protein | 43 | 201 | 518.7 | 1137 | 8.1 |
| Membrane fusion protein | 16 | 201 | 260.2 | 318 | 6.4 |
| Molecular chaperone | 18 | 219 | 347.7 | 522 | 8.9 |
| Outer membrane protein | 8 | 234 | 250.9 | 264 | 7 |
| Putisolvin related integral membrane protein | 53 | 201 | 369.2 | 783 | 6.7 |
| Putisolvin synthetase | 1 | 309 | 309 | 309 | 3.8 |
| **Mix-2** | | | | | |
| Heat-shock protein | 36 | 201 | 505.2 | 1125 | 8.4 |
| Membrane fusion protein | 10 | 225 | 269.4 | 318 | 6.7 |
| Molecular chaperone | 15 | 204 | 368.4 | 528 | 9.2 |
| Outer membrane protein | 9 | 210 | 241 | 264 | 6.4 |
| Putisolvin related integral membrane protein | 51 | 201 | 370.1 | 777 | 6.2 |
| Putisolvin synthetase | 1 | 333 | 333 | 333 | 4.4 |
| **Orange-1** | | | | | |
| Heat-shock protein | 39 | 204 | 596.2 | 1155 | 7 |
| Membrane fusion protein | 13 | 225 | 269.5 | 321 | 5.3 |
| Molecular chaperone | 20 | 237 | 360.4 | 528 | 7.2 |
| Outer membrane protein | 16 | 201 | 232.5 | 264 | 6.3 |
| Putisolvin related integral membrane protein | 57 | 201 | 360.5 | 777 | 5.4 |
| Putisolvin synthetase | 3 | 213 | 258 | 333 | 3.4 |
| **Orange-2** | | | | | |
| Heat-shock protein | 61 | 201 | 564.6 | 1137 | 6.6 |
| Membrane fusion protein | 21 | 216 | 259.1 | 327 | 5.2 |
| Molecular chaperone | 26 | 225 | 376.7 | 528 | 7.2 |
| Outer membrane protein | 22 | 204 | 255.4 | 519 | 5.5 |
| Putisolvin related integral membrane protein | 65 | 201 | 408 | 777 | 5.8 |
| Putisolvin synthetase | 3 | 207 | 270 | 333 | 4.1 |

Table 2: Basic statistics for the gene-centric assembly of surfactant-related genes. For one gene involved in iturin A biosynthesis, one in lichenysin biosynthesis and two in mycosubtilin biosynthesis, we report the number of contigs, their minimum, mean and maximum length and their average coverage, based on the separate assembly of all six samples.

| Gene name | Number of contigs | Min. | Mean Length | Max. | Average coverage |
|---|---|---|---|---|---|
| **Mango-1** | | | | | |
| Iturin A synthetase C | 1 | 213 | 213 | 213 | 4.9 |
| Lichenysin synthetase A | 8 | 246 | 310.1 | 429 | 7.3 |
| mycosubtilin synthase subunit A | 4 | 201 | 268.5 | 450 | 5.9 |
| Mycosubtilin synthase subunit B | 3 | 201 | 242 | 309 | 4.1 |
| **Mango-2** | | | | | |
| Iturin A synthetase C | 1 | 207 | 207 | 207 | 4.3 |
| Lichenysin synthetase A | 8 | 246 | 325.1 | 438 | 9.1 |
| mycosubtilin synthase subunit A | 1 | 450 | 450 | 450 | 6.9 |
| Mycosubtilin synthase subunit B | 5 | 216 | 262.8 | 345 | 4.2 |
| **Mix-1** | | | | | |
| Iturin A synthetase C | 1 | 216 | 216 | 216 | 7.6 |
| Lichenysin synthetase A | 4 | 315 | 326.2 | 336 | 5.8 |
| mycosubtilin synthase subunit A | 3 | 204 | 225 | 258 | 3.3 |
| Mycosubtilin synthase subunit B | 1 | 216 | 216 | 216 | 4.7 |
| **Mix-2** | | | | | |
| Iturin A synthetase C | 2 | 216 | 216 | 216 | 7.2 |
| Lichenysin synthetase A | 2 | 318 | 327 | 336 | 5.9 |
| mycosubtilin synthase subunit A | 2 | 204 | 208.5 | 213 | 3.5 |
| Mycosubtilin synthase subunit B | 1 | 216 | 216 | 216 | 6.9 |
| **Orange-1** | | | | | |
| Iturin A synthetase C | 3 | 204 | 207 | 213 | 2.6 |
| Lichenysin synthetase A | 12 | 219 | 306 | 444 | 6.4 |
| mycosubtilin synthase subunit A | 21 | 204 | 780.4 | 2820 | 3.3 |
| Mycosubtilin synthase subunit B | 13 | 201 | 250.2 | 276 | 3 |
| **Orange-2** | | | | | |
| Iturin A synthetase C | 1 | 207 | 207 | 207 | 2.1 |
| Lichenysin synthetase A | 6 | 246 | 315 | 444 | 9.4 |
| mycosubtilin synthase subunit A | 2 | 204 | 327 | 450 | 5 |
| Mycosubtilin synthase subunit B | 1 | 306 | 306 | 306 | 4.3 |
